# Rapid recycling of glutamate transporters on the astroglial surface

**DOI:** 10.1101/2020.11.08.373233

**Authors:** Piotr Michaluk, Janosch Heller, Dmitri A. Rusakov

## Abstract

Glutamate uptake by high-affinity astroglial transporters confines excitatory transmission to the synaptic cleft. The efficiency of this mechanism depends on the transporter dynamics in the astrocyte membrane, which remains poorly understood. Here, we visualise the main glial glutamate transporter GLT1 by generating its functional pH-sensitive fluorescent analogue, GLT1-SEP. Combining FRAP-based methods with molecular dissection shows that 70-75% of GLT1-SEP are expressed on the astroglial surface, recycling with a lifetime of only ~22 s. Genetic deletion of the C-terminus accelerates GLT1-SEP membrane turnover by ~60% while disrupting its molecule-resolution surface pattern as revealed by dSTORM. Excitatory activity boosts surface mobility of GLT1-SEP, involving its C-terminus, metabotropic glutamate receptor activation, intracellular Ca^2+^ signalling and calcineurin-phosphatase activity, but not the broad-range kinase activity. The results suggest that membrane turnover, rather than than lateral diffusion, is the main ‘redeployment’ route for the immobile fraction (20-30%) of surface-expressed GLT1. This reveals a novel mechanism by which the brain controls extrasynaptic glutamate escape, in health and disease.

## INTRODUCTION

Excitatory transmission in the brain occurs mainly through the release of glutamate at chemical synapses. Once released, glutamate is taken up by high-affinity transporters that densely populate the plasma membrane of brain astrocytes (Wadiche et al., 1995a; Danbolt, 2001). The main glial glutamate transporter GLT1 (EAAT2) maintains extracellular glutamate at nanomolar levels, thus constraining its excitatory action mainly to the synaptic cleft (Moussawi et al., 2011; Zheng and Rusakov, 2015). Because synaptic vesicles release ~3000 glutamate molecules (Savtchenko et al., 2013) and because glutamate uptake cycle can take tens of milliseconds (Wadiche et al., 1995b), large numbers of transporter molecules have to be available near synapses to buffer the escaping glutamate (Lehre and Danbolt, 1998; Bergles et al., 2002). Indeed, the high occurrence of GLT1 in astroglial plasma membranes (Danbolt, 2001) ensures that regular network activity does not overwhelm glutamate transport (Bergles and Jahr, 1998; Diamond and Jahr, 2000). However, intense excitation can prompt glutamate escape from the immediate synapse, leading to activation of extrasynaptic receptors or even neighbouring synapses (Lozovaya et al., 1999; Arnth-Jensen et al., 2002; Scimemi et al., 2004). Ultimately, the reduced availability of GLT1 has long been associated with pathologic conditions such as neurodegenerative diseases, epilepsy, or stroke (Maragakis and Rothstein, 2004; Fontana, 2015).

These considerations prompted intense interest in the cellular mechanisms underlying cellular trafficking and turnover of astroglial and neuronal glutamate transporters. A growing body of evidence has suggested the involvement of its carboxyl-terminal domain and protein kinase C (Kalandadze et al., 2002; Gonzalez et al., 2007) and calmodulin-dependent protein kinase (Underhill et al., 2015), also engaging ubiquitin-dependent processes (Gonzalez et al., 2007; Gonzalez-Gonzalez et al., 2008; Martinez-Villarreal et al., 2012) and constitutive protein sumoylation (Garcia-Tardon et al., 2012; Foran et al., 2014; Piniella et al., 2018). Ultimately, these findings unveil the potential to regulate long-term, systemic changes in the GLT1 expression in a therapeutic context (reviewed in (Fontana, 2015; Peterson and Binder, 2019)). However, what happens to GLT1 trafficking on the time scale of the ongoing brain activity remains poorly understood. In recent elegant studies, single-particle tracking with quantum dots (QDs) has detected high surface mobility of GLT1 in astroglia (Murphy-Royal et al., 2015; Al Awabdh et al., 2016). Lateral diffusivity of transporters was boosted by local glutamatergic activity, thus suggesting the use-dependent surface supply of GLT1 towards active synapses (Murphy-Royal et al., 2015; Al Awabdh et al., 2016). However, synthetic QDs almost certainly prevent their link-labelled molecules from the membrane-intracellular compartment turnover and, at the same time, do not label any newly appearing molecules on the cell surface. Thus, the molecule-tracking observations relying solely on QDs could miss important changes in the composition and/or mobility of the studied molecular species due to their continuous recycling in the membrane.

We therefore set out to develop an approach enabling us to document, in real time, the exchange between membrane and intracellular fractions of GLT1, in addition to monitoring its lateral diffusion on the cell surface. To achieve this, we generated a fully functional variant of GLT1, termed GLT1-SEP, by adding an extracellular fragment with the pH-sensitive, Super-Ecliptic pHluorin (SEP); GLT1-SEP fluoresces when exposed to the extracellular but not in low pH of intracellular compartments. Expressing GLT1-SEP in astroglia in cell cultures and brain slices allowed us to combine the optical protocols of fluorescence recovery after photobleaching (FRAP) with molecular and pharmacological dissection, to monitor membrane turnover and lateral diffusion of the transporter proteins.

## RESULTS

### Developing and probing GLT1-SEP

First, we designed the GLT1-SEP probe for FRAP measurements by introducing SEP into the second intracellular loop of GLT1a, between two proline residues (P199 and P200). Next, aiming at astrocyte-specific expression we cloned the construct under the gfaABC1D promoter (Lee et al., 2008) (Figure 1A; Methods).

**Figure 1.**
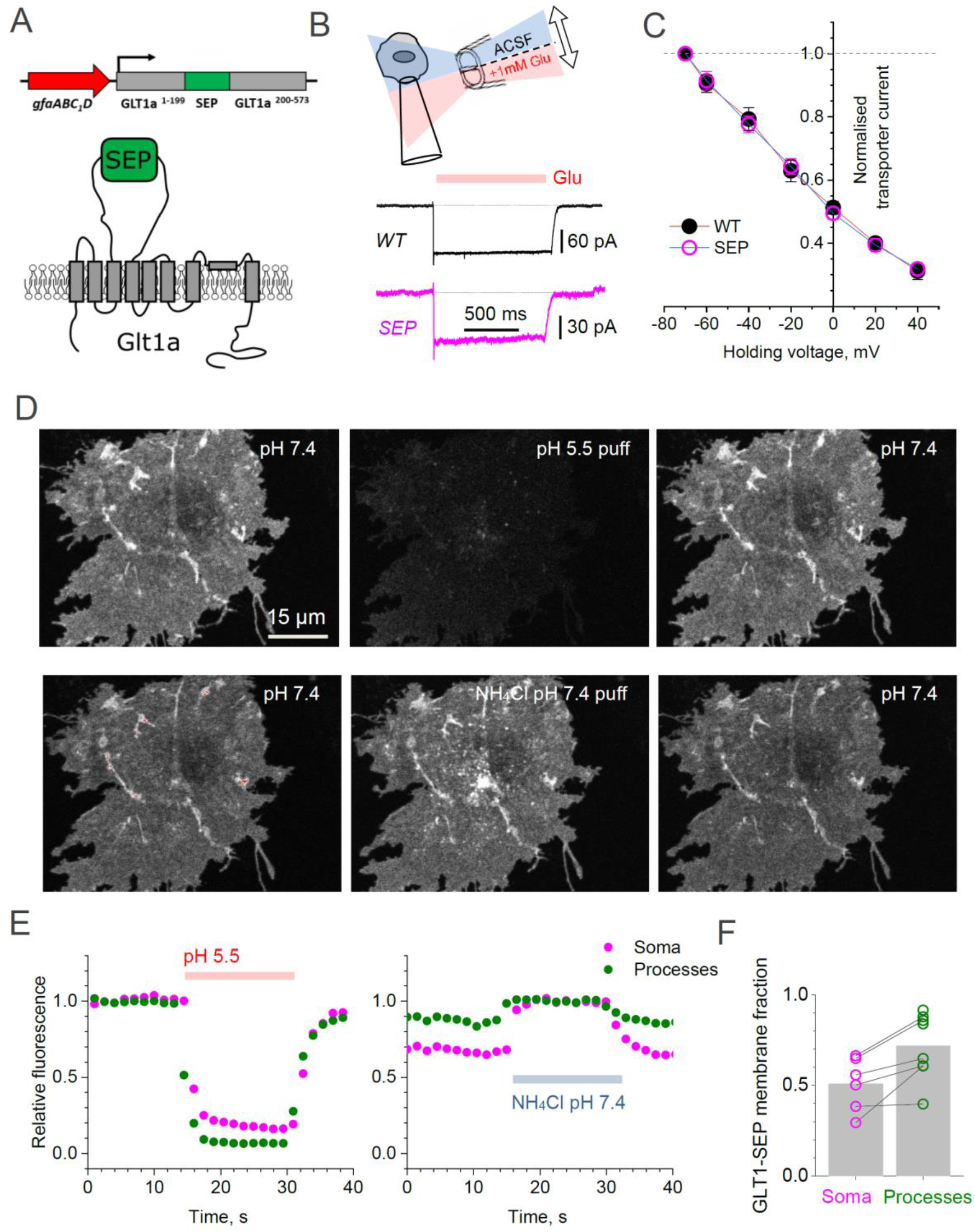
Superecliptic synaptophluorin GLT1-SEP enables monitoring of cell membrane and cytosolic fractions of glial glutamate transporters. (A) *Left:* Diagram illustrating molecular composition of GLT1-SEP. (B) Functional probing of wild-type GLT1-SEP (SEP) probe expressed in HEK cells shows a prominent current response to glutamate application, similar to that wild-type GLT-1 (WT); top diagram, theta-glass pressure pipette application; traces, one-cell examples (V_h_ = −70 mV). (C) Summary of tests shown in (A): normalised current-voltage dependencies of GLT-1 (mean ± SEM; n = 8) and GLT1-SEP (n = 4) are indistinguishable; current values normalised at V_h_ = −70 mV (absolute values 194 ± 29 pA and 81 ± 10 pA for GLT-1 and GLT1-SEP, respectively). (D) Transient acidification (~10 s pH 5.5 puff, upper row) supresses cell-surface GLT1-SEP fluorescence whereas transient membrane NH_4_^+^ permeation (~10 s NH_4_Cl puff, lower row) reveals the cytosolic fraction of GLT1-SEP; one-cell example. (E) Time course of fluorescence intensity averaged over the cell soma (magenta) or all processes (green) in the test shown in (C). (F) Average cell-surface fraction *R* of GLT1-SEP (summary of experiments shown in D-E); dots, individual cells (connecting lines indicate the same cell); grey bars, average values (*R* mean ± SEM: 0.51 ± 0.15, n = 6 for somata; 0.72 ± 0.18, n = 8 for processes; soma boundaries in two cells were poorly defined).

To test if this mutant (termed GLT1-SEP thereafter) is a functional glutamate transporter we transfected HEK 293T cells with the prepared construct. The control group of cells was transfected with the plasmid coding wild-type GLT1. For identification purposes, and to keep the same plasmid concentrations, cells were co-transfected with GLT1 constructs and mRFP1 under β-actin promoter, at a 2:1 ratio. Next, in whole-cell mode we recorded uptake currents in transfected cells induced by a 1 s application of 1 mM glutamate through a theta-glass solution-exchange system (Figure 1B), the method that avoids any mechanical concomitants of the application protocol (Sylantyev and Rusakov, 2013). Systematic recording across holding voltages produced normalized I-V curves that showed an excellent match between the wild-type native transporter and the mutant (Figure 1C). However, the absolute current in GLT1-SEP expressing cells was on average ~50% lower (Figure 1 - figure supplement 1A), possibly because of lower expression compared to native GLT1.

**Figure 1 - figure supplement 1.**
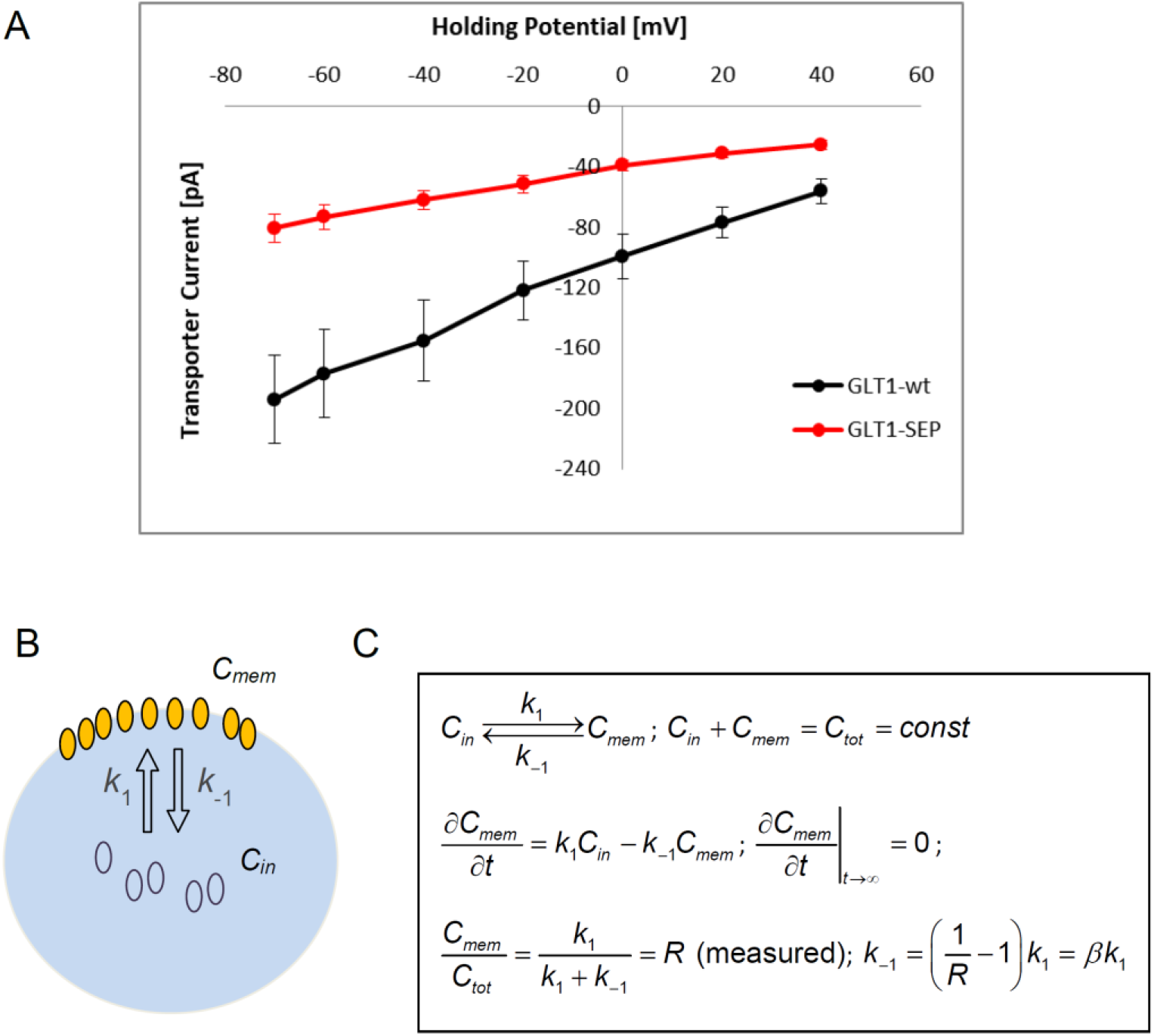
Testing glutamate transport function and the membrane/ cytosol fraction ratio for GLT1-SEP. (A) Summary of transporter current recordings in wild-type GLT-1 and GLT1-SEP expressing HEK cells, as indicated; absolute current values are shown. (B) *D*iagram illustrating the kinetics of exchange between the plasma membrane fraction (concentration *C_m_*) and the cytosol fraction (concentration *C_in_*) of GLT1-SEP; *k*_1_ and *k*_-1_, kinetic constants, as shown. (C) *Right*, kinetic equations describing membrane-cytosol exchange for GLT1-SEP; *C_tot_*, total concentration of GLT1-SEP; *R*, is the (equilibrated) membrane fraction of GLT1-SEP, measured experimentally (Figure 1).

### Intracellular versus membrane fractions of GLT1-SEP in astroglia

We next expressed GLT1-SEP in mixed cultures of neurons and glial cells. Thanks to the gfaABC1D promoter, the probe was almost exclusively expressed in astrocytes. The living GLT1-SEP expressing cells were readily visualised, featuring a dense and homogenous expression pattern that reveals fine detail of cell morphology (Figure 1D, upper left). Because the pH-sensitive GLT1-SEP fluoresces at higher extracellular pH but not at lower intracellular pH, we were able to estimate directly its membrane and intracellular fractions. Firstly, we confirmed that the observed fluorescence comes mainly from the membrane fraction of GLT-SEP. Indeed, brief acidification of the extracellular medium to pH 5.5 (10 second pipette puff) reversibly suppressed GLT1-SEP fluorescence (Figure 1D, upper row). Conversely, proton permeation of the cell membrane (10 second puff with NH_4_Cl) could reveal both intra- and extracellular GLT1-SEP fractions, in a reversible fashion (Figure 1D, lower row). Systematic quantification of these experiments (Figure 1E) provided an estimate of the average GLT1-SEP surface fraction in astroglial processes, *R* = 0.72 ± 0.18 (n = 8 cells, Figure 1F). In other words, between 2/3 and 3/4 of all cellular GLT1-SEP were exposed to the extracellular space. The *R* estimate for the cell soma was somewhat lower (Figure 1F), but because exact identification of the somatic boundaries was ambiguous, we did not use somatic data in further analyses.

These data provided an important constraint for a (steady-state) quantitative assessment of the GLT1-SEP turnover kinetics. Introducing the membrane-intracellular exchange reaction (Figure 1-figure supplement 1B) as 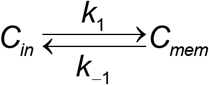 (*C_m_* and *C_in_* are membrane and intracellular concentration of GLT1-SEP, respectively) leads to a direct relationship between the corresponding kinetic constants *k*_1_ and *k*_-1_ (Figure 1-figure supplement 1C): 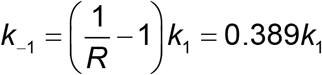. However, this steady-state relationship alone could not reveal the actual rate of GLT1-SEP turnover in the cell membrane. To address this, we implemented a different approach.

### GLT1-SEP recycling in the plasma membrane

Because photobleaching quenches irreversibly only the fluorophores that are in the excited (fluorescent) state, it could be used to separate the fluorescent from the non-fluorescent GLT1-SEP fraction. We therefore implemented a two-photon excitation FRAP protocol in which photobleaching applies virtually to the entire astrocyte expressing GLT1-SEP (Figure 2A). This was feasible mainly because the morphology of cultured astroglia was essentially two-dimensional, thus permitting comprehensive photobleaching in close proximity of the focal plane. Thus, a brief (2 s) laser scan could almost entirely suppress GLT1-SEP fluorescence within the target area (Figure 2A, dashed red circle) enabling us to document partial fluorescence recovery within smaller ROIs inside the bleached area: sampling normally included three ~10 μm wide circular ROIs over cell processes (and additionally one ~20 μm ROI over the soma) in each cell (Figure 2A, dotted orange circles). The ROI selection was restricted to morphologically homogenous cell areas inside the fully bleached territory, but otherwise was quasi-random (three ROIs picked randomly out of 10-20 available per cell).

**Figure 2.**
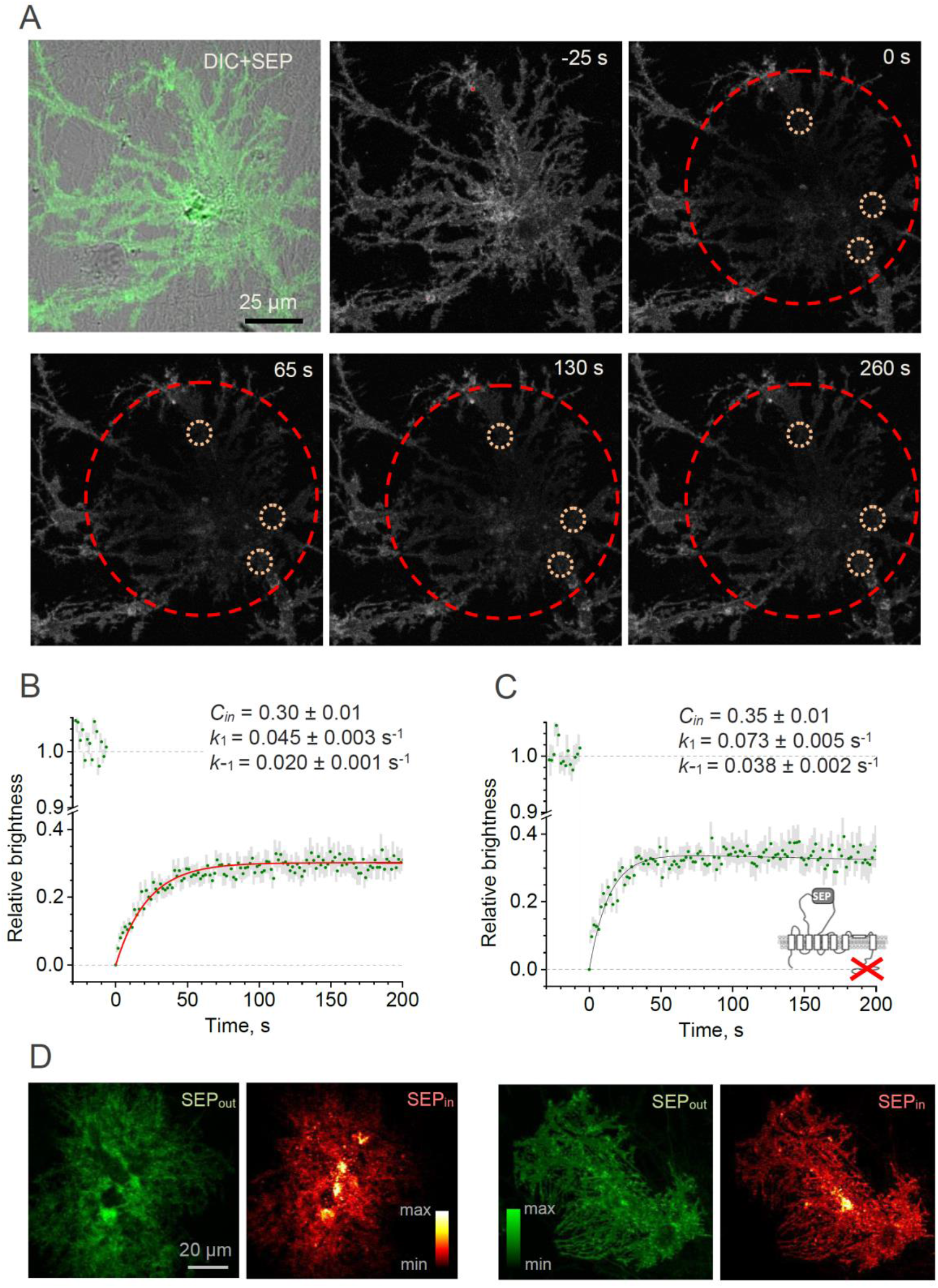
Whole-cell FRAP reveals the kinetics of the GLT1-SEP membrane surface turnover. (A) One-cell example illustrating FRAP protocol; upper left, DIC+SEP channel image; serial images, GLT 1-SEP channel at different time points (indicated) after a photobleaching pulse (*t* = 0 s); dashed red circle, laser-photobleached region; dotted orange circles, example of ROIs. (B) Time course (mean ± SEM, n = 27 ROIs in N = 9 cells) of the GLT1-SEP fluorescence intensity within the photobleached region (as in A), normalised against the baseline value. Red line, best-fit GLT1-SEP FRAP kinetics incorporating cytosolic protein fraction (*C_in_*), membrane-surface turnover constants (*k*_1_ and *k*_-1_) and the residual photobleaching constant (*k_b_*; not shown); see text and Figure 2S for further detail. (C) Experiment as in (B), but with the with the C-terminus deleted mutant GLT1ΔC-SEP expressed in astroglia (n = 25 ROIs in N = 8 cells); other notations as in (B). (D) Two characteristic examples illustrating cellular distribution of surface-bound fraction of GLT1-SEP (green, SEP_out_) and its intracellular fraction (red, SEP_in_) in live individual astroglia.

These experiments produced the average FRAP time course, with relatively low noise (Figure 2B). The cellular biophysical mechanisms underpinning this time course combine membrane insertion of non-bleached GLT1-SEP and, if any, residual photobleaching of surface-bound GLT1-SEP. Solving the corresponding kinetic equations (Figure 2-figure supplement 1) provide the resulting fluorescence time course as 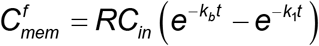 where *t* is time and *k_b_* is the residual photobleaching constant (other notations as above). This equation has two orthogonal (independent) free parameters, *C_in_* and *k_1_*, whereas the residual photobleaching rate *k_b_* turned out to be negligible throughout the sample. The best-fit estimate gave (Figure 2B): *k*_1_ = 0.045 ± 0.003 s^-1^, *k*_-1_ = 0.020 ± 0.001 s^-1^, and *C_in_* = 0.30 ± 0.01. Reassuringly, the value of *C_in_* (intracellular fraction of GLT1-SEP) obtained in these experiments was indistinguishable from the value of 1 - *R* = 0.28 obtained using a fully independent proton permeation method (Figure 1F).

**Figure 2 - figure supplement 1.**
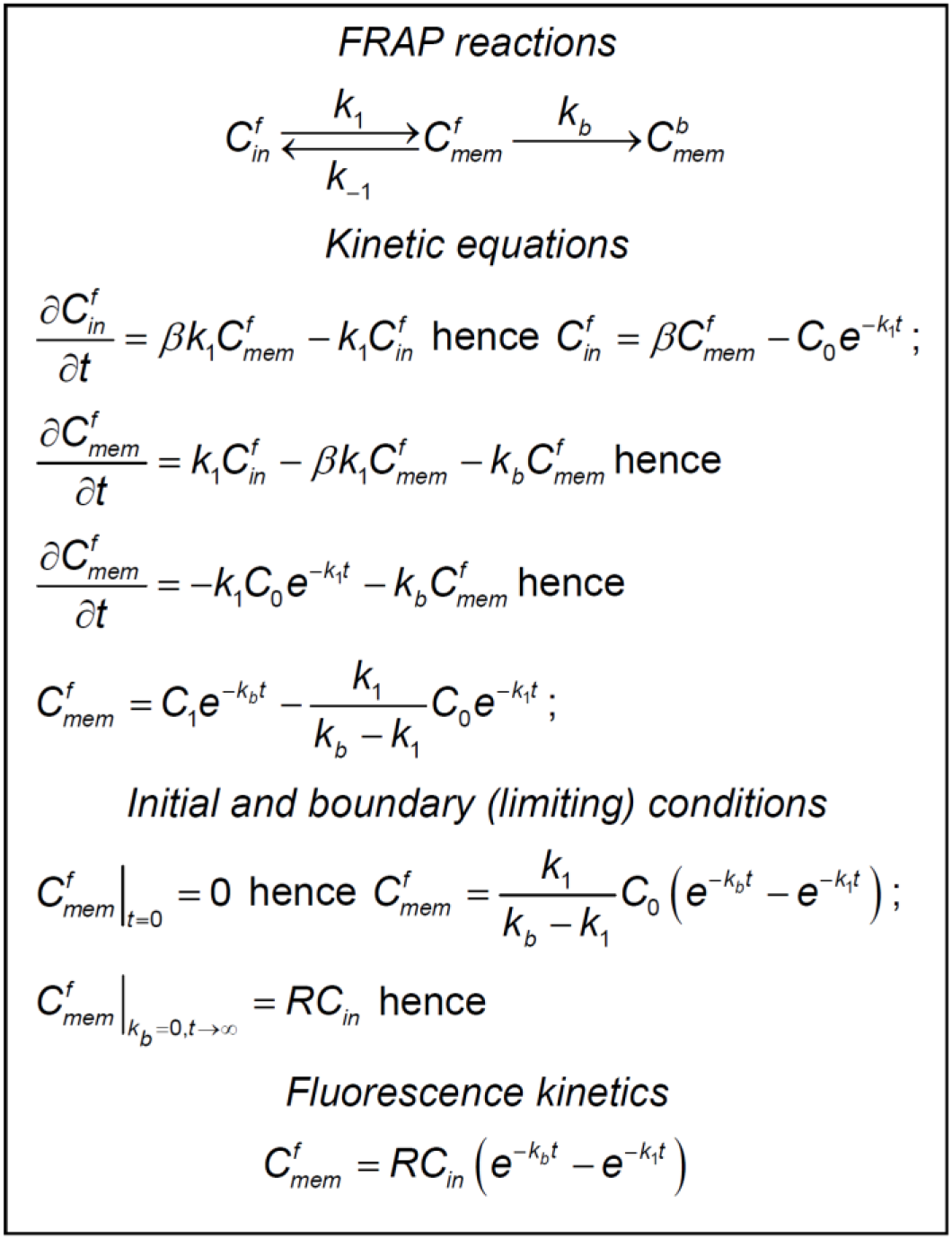
Establishing the kinetics of whole-cell FRAP for GLT1-SEP molecules in astrocytes. *FRAP reactions* diagram reflects exchange (turnover) between membrane cytosol fractions of non-bleached GLT1-SEP molecules, with 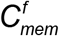 and 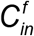 standing for their relative concentrations, respectively, and residual bleaching of the membrane fraction adding to the bleached membrane fraction 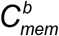. *k*_1_, *k*_-1_, and *k*_b_ are the kinetic constants, as indicated. *Kinetics equations* describe the FRAP reactions in partial derivatives for 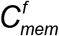 and 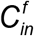. The corresponding solution includes two unknown constants, *C*_0_ and *C*_1_, which are determined using *Initial and boundary conditions*, leading to the expression of *Fluorescence kinetics*. Other notations: 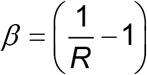 where *R* is total (bleached and non-bleached) membrane fraction of GLT1-SEP, as in Figure 1 - figure supplement 1.

These estimates suggest that the characteristic lifetime of the membrane GLT1-SEP fraction, as given by *k*_1_^−1^, is ~22 seconds. Because the cytosolic carboxy-terminal domain of GLT1 has earlier been implicated in the GLT1 expression mechanism (Gibb et al., 2007; Foran et al., 2014), we asked whether cleaving it interferes with the membrane kinetics of the transporter. We therefore generated a GLT1-SEP mutant with a deleted C-terminus, GLT1ΔC-SEP, and expressed it in astroglia. FRAP experiments in the GLT1ΔC-SEP expressing cells (Figure 2C) showed that deleting the C-terminus had only a moderate effect on the intracellular fraction of transporters (*C_in_* = 0.35 ± 0.01) but reduced the GLT1 membrane lifetime by nearly a half (to ~14 s). This finding suggests that the C-terminus could play an important role in retaining GLT1 in the plasma membrane, even though a steady-state membrane-intracellular compartment ratio remains almost unaffected.

Do the cell-average values of the GLT1-SEP membrane fraction (and hence turnover rate) occur homogeneously throughout the cell morphology? To understand this, we directly compared distributions of the membrane and the intercellular populations of GLT1-SEP: the latter was obtained by subtracting the surface GLT1-SEP image from the total GLT1-SEP image (under NH_4_Cl, as in Figure 1D). Intriguingly, this comparison revealed that the membrane GLT1-SEP does not necessarily predict the intracellular GLT1-SEP pattern which could display prominent clustering features (Figure 2D). Thus, at least in some cases the membrane dynamics of GLT1 could be specific to microscopic regions of the cell.

### Nanoscale distribution of GLT1 species with respect to synapses

While our FRAP approach measures live GLT1 turnover in the astrocyte membrane, it does not reveal surface distribution of these molecules, in particular that with respect to synaptic connections. We therefore turned to super-resolution microscopy that involves stochastic localisation of individual molecules dSTORM (van de Linde et al., 2011) using multi-colour 3D STORM experimental protocols that we have established previously (Heller et al., 2017; Heller and Rusakov, 2019; Heller et al., 2020) (Methods). We thus used chromatically separable photoswitchable dyes to visualise distributions of the wild type GLT 1, mutant GLT1-SEP, or GLT 1ΔC-SEP species and their relationship to the synaptic clusters of the ubiquitous postsynaptic density protein PSD95 in mixed cultures (Figure 3A, Figure 3-figure supplement 1A).

**Figure 3.**
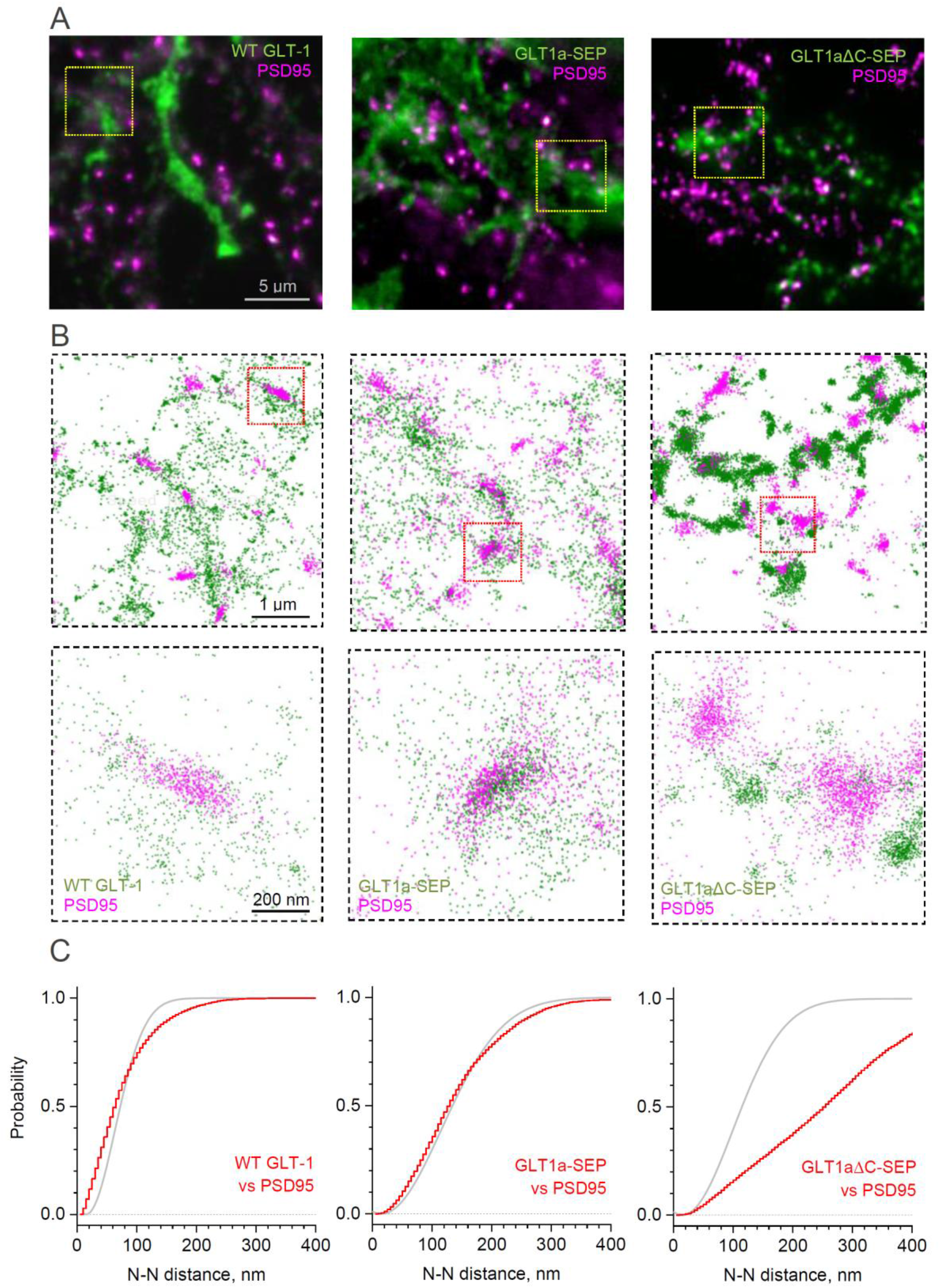
Distribution of GLT-1 species in relation to postsynaptic densities in the astroglial membrane: A super-resolution dSTORM analysis. (A) Wide-field fluorescent images (examples) illustrating antibody labelled GLT1 species (green channel) and postsynaptic density protein PSD95 (magenta), as indicated, in mixed astroglia-neuron cultures. See Figure S3A for macroscopic views. (B) dSTORM nano-localisation maps (examples) depicting individual labelled GLT1 species (green), as indicated, and PSD95 (magenta) molecules. Top row, ROIs shown as the corresponding yellow squares in (A); bottom row, ROIs shown as red squares in the top row. (C) Red line (5 nm bins): distribution *D*(*r*) of nearest-neighbour (N-N) distances *r* between labelled GLT1 species and clusters of PSD95 molecules (PSD95 clusters represent >50 particles <100 nm apart). Grey line: theoretical distribution *D*(*r*) = 1 − exp(−*λπr*^2^) that corresponds to the Poisson point process (evenly random scatter) with the same surface density of PSD95 clusters *λ* as sampled experimentally. Experimental *λ* values were: 67 μm^-2^ (WT GLT-1), 15.5 μm^-2^ (GLT1-SEP), and 22.2 μm^-2^ (GLT1Δ-SEP); see Methods for detail.

dSTORM visualisation revealed that the scatter of wild-type GLT1 tends towards forming clusters, both among GLT1 molecules and also between GLT1 and PSD95, and that GLT1-SEP-expressing cells display similar features (Figure 3B). Indeed, the classical nearest-neighbour analysis indicated that the pattern of wild-type GLT1 and GLT1-SEP with respect to PSD95 clusters deviates from the evenly random distribution towards closer spatial association (Figure 3C), and that these transporter molecules also tend to form short-distance (up to 50 nm) clusters among themselves (Figure 3-figure supplement 1B).

**Figure 3 - figure supplement 1.**
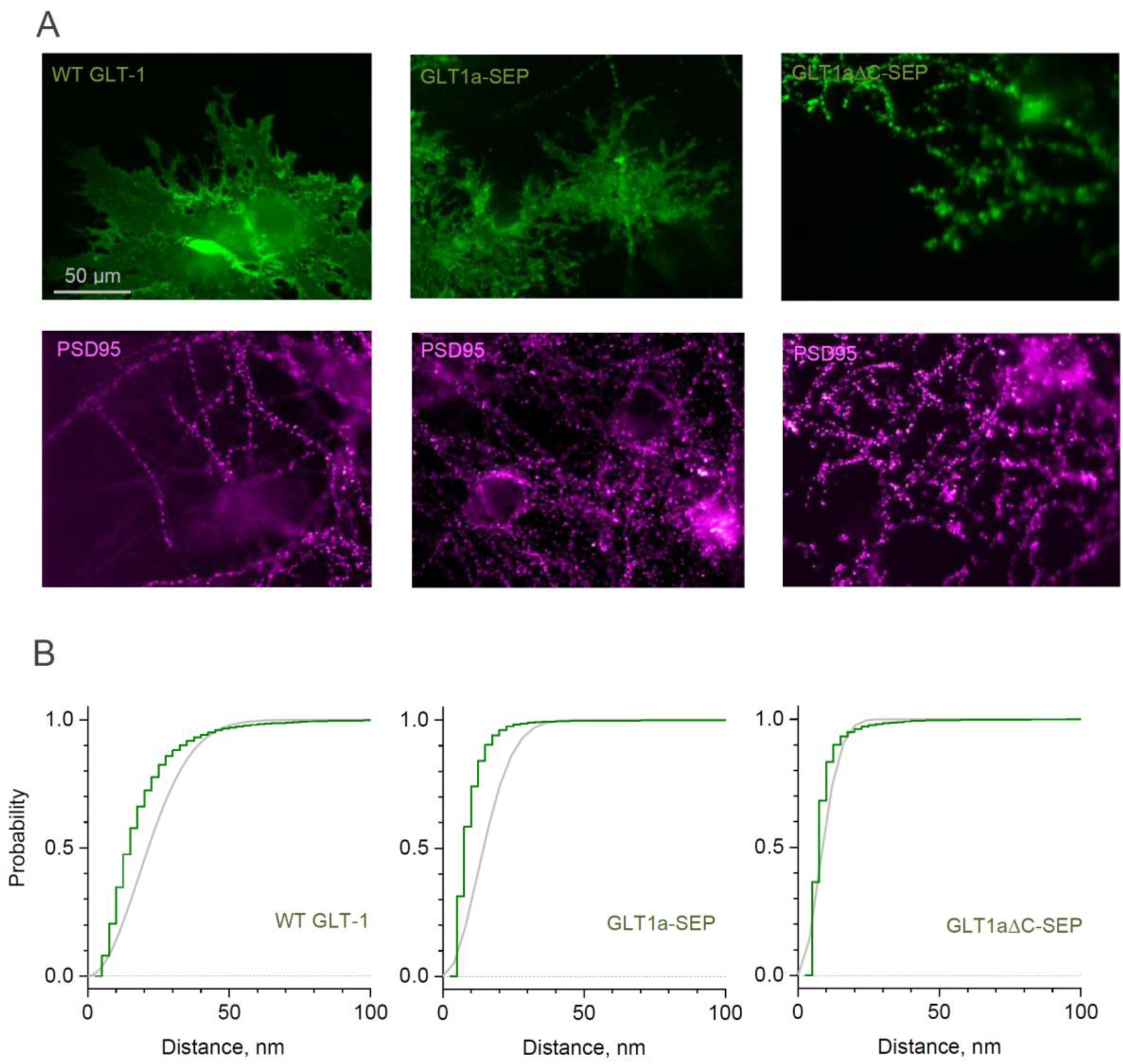
Distribution of GLT-1 species in the astroglial membrane: macroscopic wide-field view and super-resolution dSTORM analysis. (A) Wide-field fluorescent images (examples) displaying antibody labelled GLT1 species (green channel) and postsynaptic density protein PSD95 (magenta), as indicated, in mixed astroglia-neuron cultures. See Figure 3 for higher magnification. (B) dSTORM analyses (see Figure 3 for single-molecule maps): Distribution *D*(*r*) of nearest-neighbour (N-N) distances *r* among labelled GLT1 species (green line, 5 nm bins), and the theoretical distribution for the Poisson point process (evenly random scatter, *D*(*r*) = 1 – exp(−*λπr*^2^)) with the same surface density *λ* (grey line); experimentally sampled *λ* values were 324 μm^-2^ (WT GLT-1), 67 μm^-2^ (GLT1-SEP), and 2938 μm^-2^ (GLT1ΔC-SEP); a shift to the left for the red versus grey line indicates significant clustering. See Methods for detail.

In contrast, the species with deleted C-terminus, GLT1ΔC-SEP, showed spatial dissociation (distancing) with PSD95 clusters (Figure 3B-C) while displaying dense molecular clustering among themselves, to the extent that the latter is not distinguishable from uniform packing at a high local density (Figure 3-figure supplement 1B; this analysis does not cover higher-order, longer-distance GLT1ΔC-SEP clustering, which is evident in Figure 3B). These observations indicate that the C-terminus of GLT1 plays a critical role not only in its cellular membrane turnover but also in the surface expression pattern of the protein.

### Lateral mobility of GLT1-SEP in astroglia

We next set out to assess lateral surface mobility of GLT1-SEP using a classical FRAP protocol, in which the fluorescence kinetics is monitored within a small ROI (Figure 4A). Because running a FRAP protocol bleaches immobile molecules that remain within the ROI, repeating this protocol within the same ROI may produce a different FRAP time course. To account for this and any other use-dependent trends in the imaging conditions, we routinely recorded pairs of FRAP trials separated by 1 min (Figure 4B, Figure 4-figure supplement 1), unless indicated otherwise. This time interval was also longer than the GLT1 membrane turnover period (~22 s, see above), which should help minimise the number of bleached immobile molecules remaining within the ROI, as they are replaced by new arrivals from the intracellular compartment. This approach enabled us to compare FRAP kinetics between control conditions and during ligand application, in the manner that provides correction for any consistent difference within paired FRAP trials (Figure 4C, Figure 4-figure supplement 1).

**Figure 4.**
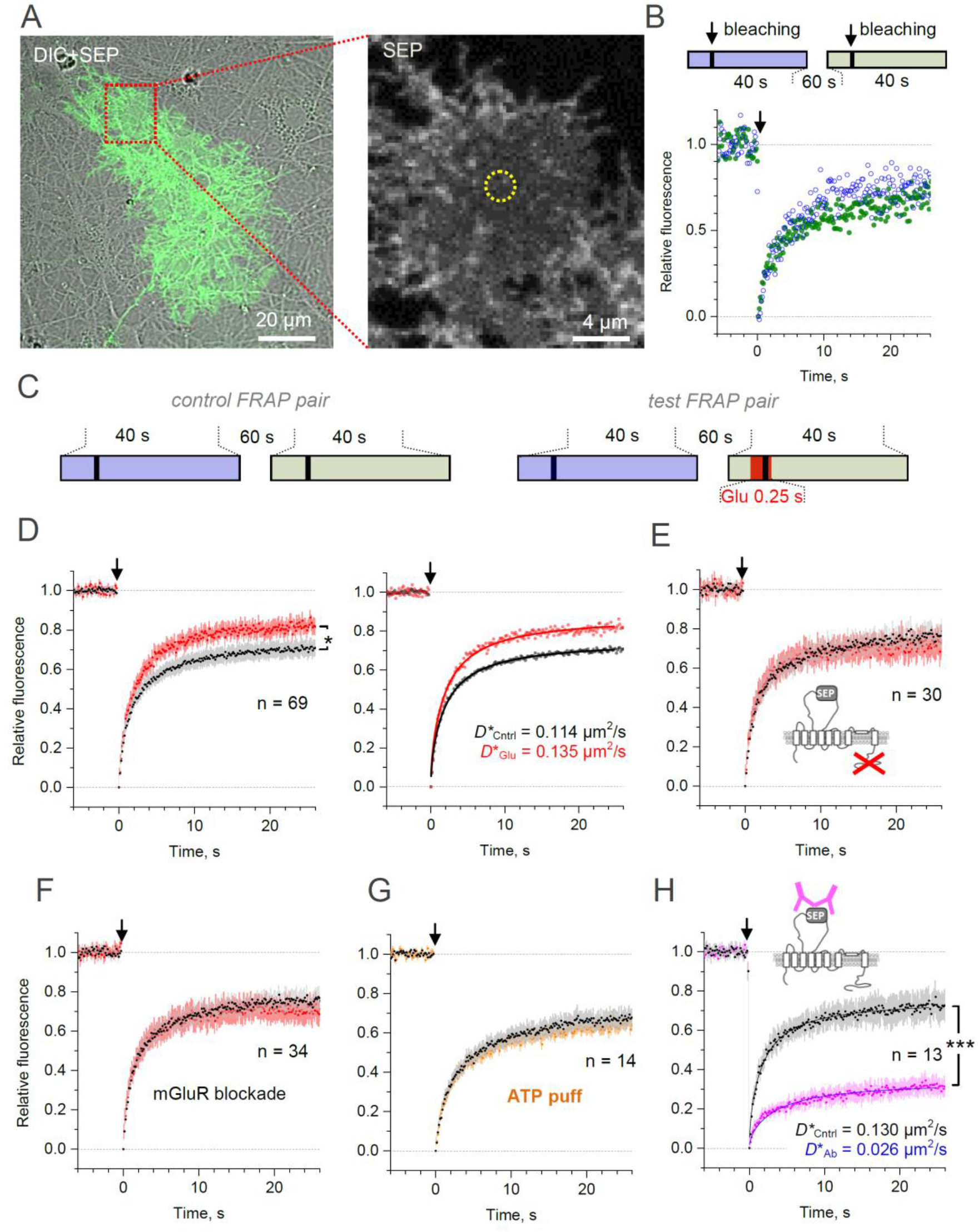
Microscopic-ROI FRAP probes lateral membrane mobility of GLT1-SEP in cultured astroglia. (A) One-cell example as seen in GLT1-SEP + DIC channel (left), with a selected area (dotted rectangle) illustrating a circular, 2.06 μm wide FRAP spot (dotted circle, right). (B) Diagram, the paired-sample FRAP protocol, in which two trials are carried out in succession, to account for any non-specific, time-dependent drift in FRAP kinetics. Plots, one-cell example of the paired-sample FRAP test, with the first and second trial data are shown in blue and green, respectively; arrow, bleaching pulse (λ_x_^2p^ = 690 nm, 10-15 mW under the objective, duration 46 ms); fluorescence ROI, photobleaching spot as in (A). (C) Diagram illustrating the paired-sample FRAP protocol, which includes both control and glutamate application cycles; FRAP kinetics under glutamate application could be corrected for non-specific drift by using the control cycle data. (D) *Left*, average time course of the GLT1-SEP FRAP (dots and shade: mean ± 95% confidence interval, here and thereafter) in baseline conditions (black) and upon glutamate application (250 ms puff 200 ms before the photobleaching pulse lured); asterisk, p < 0.05 (n = 69 FRAP spots in N = 13 cells). *Right*, FRAP time course (mean values) fitted with the Soumpasis FRAP equation for (see main text) for control and glutamate tests. Best-fit GLT1-SEP diffusion coefficient *D* is shown for control (Cntrl) and glutamate puff (Glu) trials, as indicated. (E) Average FRAP time course in control and glutamate-puff tests carried out with the C-terminus deleted mutant CLT1ΔC-SEP, as indicated (n = 30 FRAP spots in N = 7 cells); other notation as in (D). (F) Average FRAP time course in control and glutamate-puff tests in the presence of AMPA and metabotropic glutamate receptor (mGluR) blockers (n = 34 FRAP spots in N = 7 cells): MPEP (1 mM), LY341495 (30 nM), YM298198 (0.3 μM); NBQX (10 μM) was added to suppress network hyper-excitability under LY341495; other notation as in (D). (G) Average FRAP time course in control conditions and after the ATP pressure puff (100 μM, 250 ms duration 200 ms before bleaching start, no glutamate), as indicated (n = 14 FRAP spots in N = 4 cells); other notation as in (D). (H) Control test: Average FRAP time course in control conditions and under surface cross-linkage by anti-GFP antibody, as indicated (n = 13 FRAP spots in N = 2 cells); other notation as in (D).

In the first experiment, we therefore documented FRAP kinetics within a small (~1.6 μm diameter) circular membrane area of a visualised astrocyte, in baseline conditions and during a brief (250 ms, 1 mM) application of glutamate 200 ms prior to bleaching onset, to mimic a transient rise in local excitatory activity.

The data (corrected for paired-trial trends) showed a clear difference in the FRAP kinetics between the two conditions (Figure 4D, left). This could reflect a difference in lateral diffusivity of mobile transporters, but also in the immobile versus mobile fractions of GLT1-SEP. To evaluate both variables from the FRAP kinetics, we used the well-established Soumpasis approach for circular ROIs (Soumpasis, 1983; Kang et al., 2009) (Methods). This fitting method operates with only two mutually independent (orthogonal) free parameters, mobile fraction *C_mob_* and diffusion coefficient *D*, and its estimates should not depend on residual changes in fluorescence, such as photobleaching (Soumpasis, 1983; Kang et al., 2009).

**Figure 4 - figure supplement 1.**
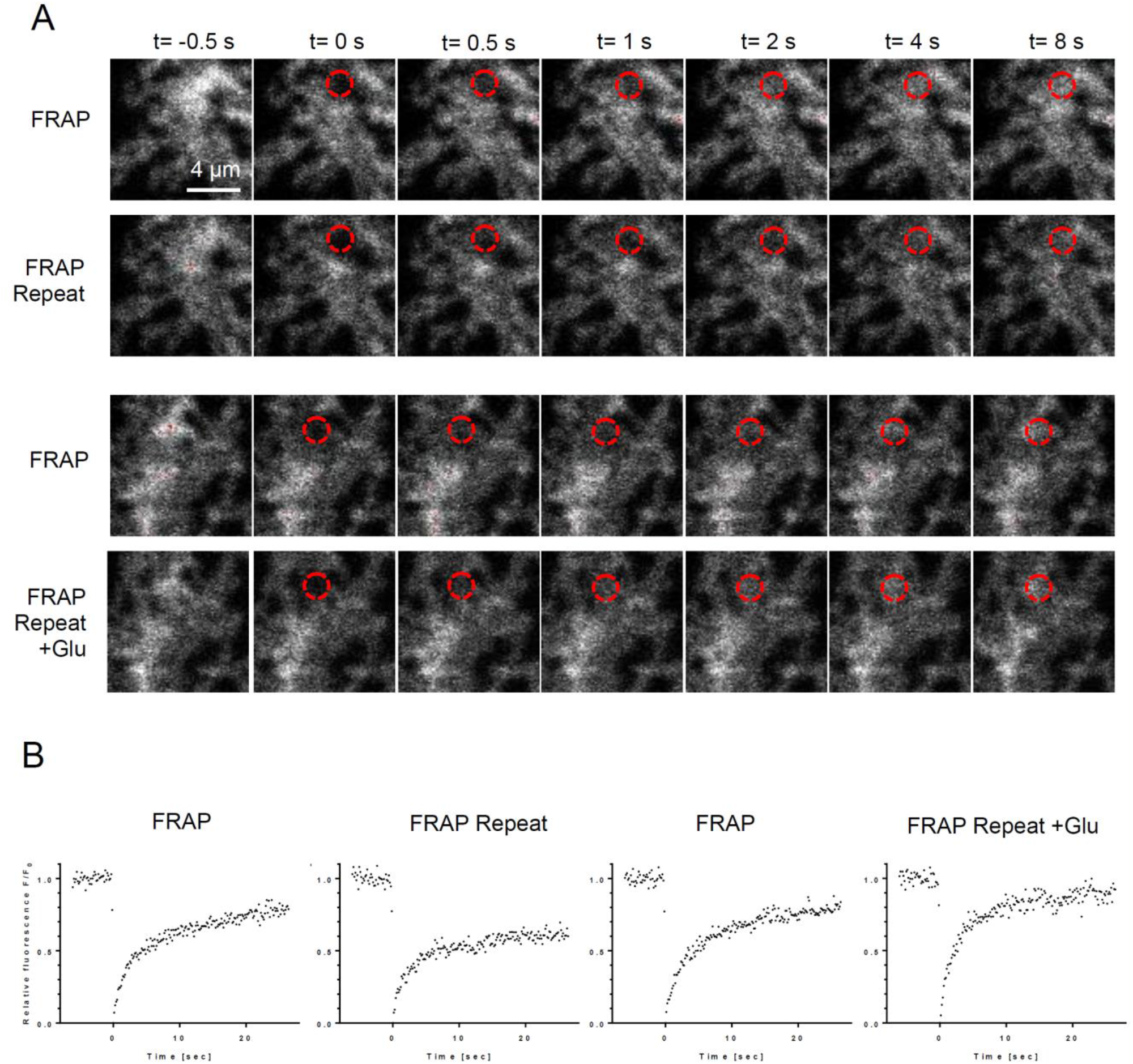
Microscopic-ROI FRAP probes lateral membrane mobility of GLT1-SEP in cultured astroglia. (A) One-cell example of FRAP kinetics at four trials (two paired-trial FRAP stages); time-lapse images of a cell fragment with a FRAP spot (dotted red circle), at selected time points before and after the photobleaching pules (at t = 0), as indicated. (B) Time course of FRAP for the four consecutive trials shown in (A), as indicated.

In baseline conditions, the best-fit values were *C_mob_* = 0.76 ± 0.01 and mobile-fraction diffusivity *D* = 0.152 μm^2^/s (diffusion time *τ_D_* = 1.75 ± 0.03, Methods), thus giving the average diffusivity (accounting for mobile and immobile molecules) *D*\* = *C_mob_ · D* = 0.114 μm^2^/s (Figure 4D, right). This value appears in correspondence with the average lateral diffusivity of GLT1 measured earlier with quantum dots (Murphy-Royal et al., 2015), although it is higher than the values reported using a different QD approach (Al Awabdh et al., 2016). When glutamate was briefly applied immediately before and after the photobleaching pulse, diffusivity of the mobile-fraction only did not appear to be affected (*τ_D_* = 1.73 ± 0.03) whereas its size has increased significantly (*C_mob_* = 0.88 ± 0.003), giving average *D*\* = 0.135 μm^2^/s, an increase of ~18% compared to control (Figure 4D, right). This result suggested that glutamatergic activity could boost overall membrane mobility of GLT1 transporters, a conclusion similar to that drawn earlier using QDs (Murphy-Royal et al., 2015; Al Awabdh et al., 2016).

### Molecular regulators of activity-dependent membrane mobility of GLT1

We next found that deleting the C-terminus of GLT1-SEP does not alter its mobility in basal conditions (*C_mob_* = 0.815 ± 0.003; *D*\* = 0.117 μm^2^/s) but appears to block the mobility-boosting effect of glutamate application (Figure 4E). A similar result was obtained when metabotropic glutamate receptors were blocked by a pharmacological cocktail: no detectable effect on GLT1-SEP mobility in baseline conditions (*C_mob_* = 0.795 ± 0.003; *D*\* = 0.121 μm^2^/s) but suppression of the glutamate-induced mobility increase (Figure 4F). Because purinergic receptors mediate a major signalling cascade in brain astroglia (Verkhratsky and Nedergaard, 2018), we asked whether ATP application alters mobility of GLT1-SEP, and detected no effect (Figure 4G).

Finally, to assess sensitivity and the dynamic range of our FRAP protocol we cross-linked surface GLT-SEP, by incubating cultures briefly (10 min in humidified incubator) with either IgY antibody (100 μg/ml, chicken polyclonal, Merck AC146) or with the anti-GFP antibody (100 μg/ml, chicken polyclonal, Abcam ab13970). The cross-linkage reduced the FRAP-measured transporter mobility five-fold (Figure 4H), confirming high sensitivity and general suitability of the present FRAP method.

### Cellular mechanisms affecting GLT1 mobility in hippocampal slices

**Figure 5 - figure supplement 1.**
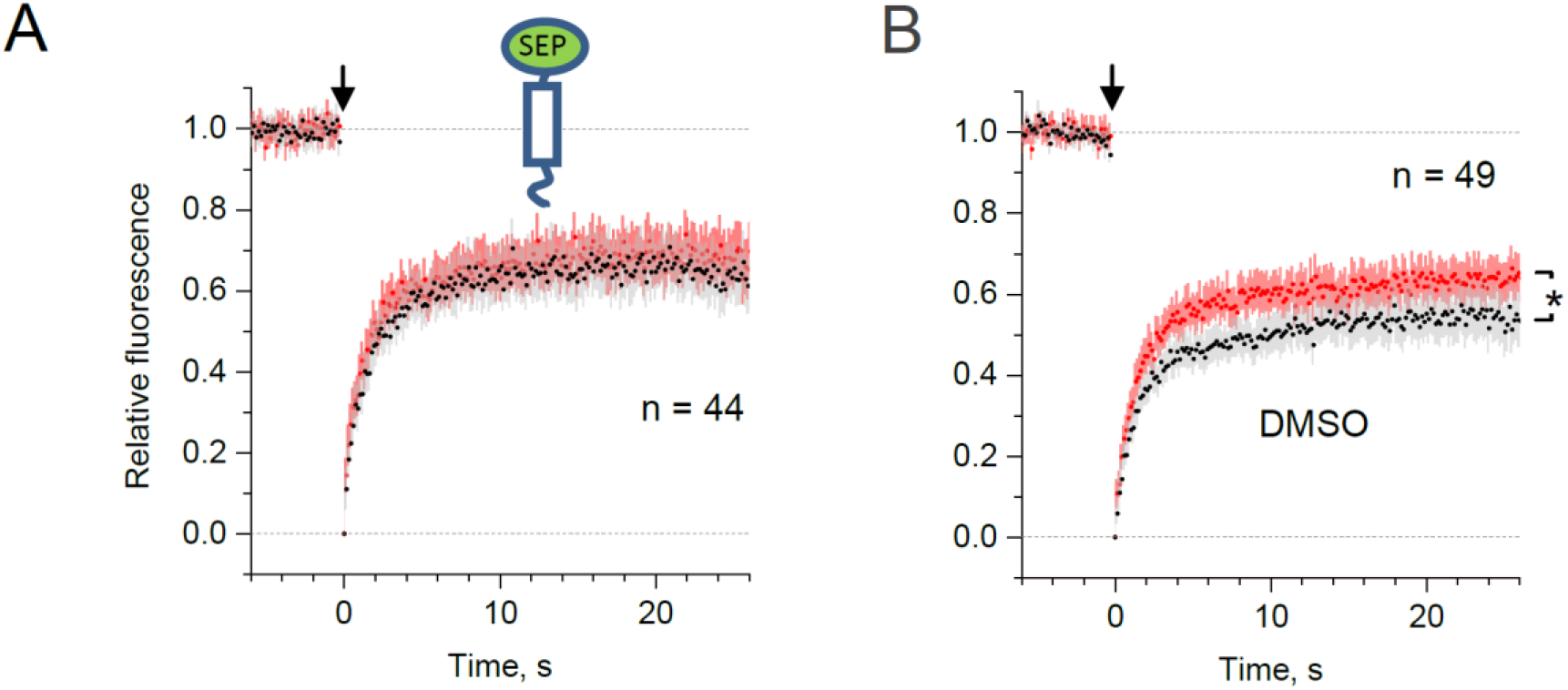
Control tests for microscopic-ROI FRAP probing of lateral membrane mobility of GLT1-SEP in organotypic hippocampal slices. (A) Average time course of the truncated transmembrane protein (C-terminal transmembrane anchoring domain of platelet-derived growth factor receptor) in fusion with SEP (dots and shade: mean ± 95% confidence interval, here and thereafter), in baseline conditions (black) and after the Bicuculine+4-AP application (red; n = 44 FRAP spots in N = 11 cells). (B) Average FRAP time course in the presence of drugs vehicle – 0.2% DMSO baseline conditions (black) and after the Bicuculine+4-AP application (red); *, p < 0.05 (n = 49 FRAP spots in N = 13 cells); other notation as in (A).

Whilst cultured astroglia are thought to retain key molecular mechanisms acting in situ, astrocytes in organised brain tissue have distinct morphology and engage in network signalling exchange that may be different from cultures. We therefore set out to validate our key observations focusing on area CA1 astroglia in organotypic hippocampal slices: these cells closely resemble their counterparts in vivo (Figure 5A), and are embedded in a well-defined synaptic circuitry. To induce a rapid rise in the spontaneous excitatory activity of the native network, we blocked GABA_A_ receptors and potassium channels with bicuculline and 4-AP (rather than applying glutamate, Figure 5B).

**Figure 5.**
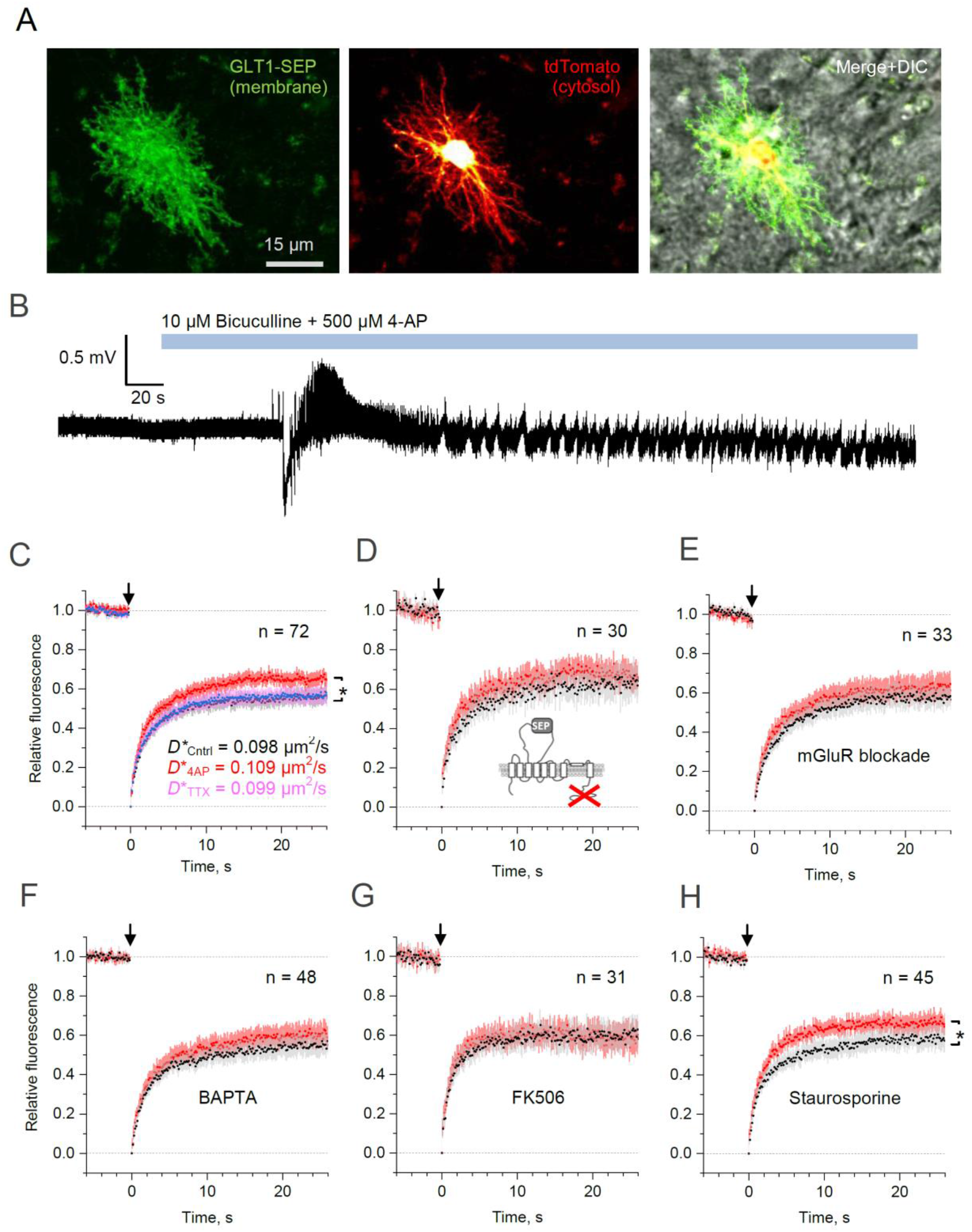
Microscopic-ROI FRAP probes lateral membrane mobility of astroglial GLT1-SEP in organotypic hippocampal slices. (A) Example of astroglia in an organotypic slice, seen in GLT1-SEP, tdTomato, and merge+DIC channel, as indicated. (B) One-slice example of boosted excitatory activity (field potential recording, CA1 area) induced by the application of GABA_A_ receptor blocker Bicuculine and the potassium channel blocker 4-AP, as indicated. (C) Average time course of the GLT1-SEP FRAP (dots and shade: mean ± 95% confidence interval, here and thereafter) in baseline conditions (black), Bicuculine+4-AP application (red), and after sodium channel blockade by TTX (magenta), as indicated; p < 0.05 (n = 72 FRAP spots in N = 15 cells). *Right*, FRAP time course (mean values) fitted with the Soumpasis FRAP equation for (see main text) for control and glutamate tests. Best-fit GLT1a-SEP diffusion coefficient *D*\* (Soumpasis FRAP fit) is shown for control (Cntrl), Bicuculine+4-AP application (4AP) and TTX trials, as indicated. (D) Average FRAP time course for the C-terminus deleted mutant GLT1aΔC-SEP, as indicated; other notation as in (C). (E) Average FRAP time course in the presence of metabotropic glutamate receptor blockers (n = 33 FRAP spots in N = 8 cells): MPEP (1 μM), LY341495 (30 nM), YM298198 (0.3 μM), and NBQX (10 μM); other notation as in (C). (F) Average FRAP time course in the presence of intracellular BAPTA (n = 48 FRAP spots in N = 10 cells); other notation as in (C). (G) Average FRAP time course under the calcineurin (phosphatase) blockade by FK506 (1 μM; n = 31 FRAP spots in N = 6 cells); other notation as in (C). (H) Average FRAP time course in the presence of the broad-range kinase activity blocker Staurospotine (100 nM); *p < 0.05 (n = 45 FRAP spots in N = 8 cells) other notation as in (C).

Because the morphology of astroglia in brain tissue is essentially three-dimensional, the whole-cell FRAP protocols (as in Figure 2A) were not technically feasible. However, the small-ROI FRAP experiments (as in Figure 4A) in slices showed that diffusivity of GLT1-SEP in the plasma membrane was similar, if somewhat slower, than that in cultures (Figure 5C). Similar to the case of cultured astroglia, elevated excitatory activity increased GLT1-SEP mobility, which could be reversed by blocking spiking activity with TTX (Figure 5C). We confirmed that this effect was not due to some unknown concomitants of increased network activity that might affect astrocyte membrane properties per se: a truncated sham-protein probe carrying an extracellular SEP domain showed no changes in lateral diffusion under this protocol (Figure 5 - figure supplement 1A). Conversely, application of the vehicle DMSO on its own had no effect on the activity-dependent increase in GLT1-SEP diffusion (Figure 5-figure supplement 1B).

Again, deletion of the C-terminus or the pharmacological blockade of metabotropic glutamate receptors suppressed the activity-dependent mobility increase (Figure 5D-E). Because metabotropic glutamate receptors engage a major Ca^2+^ signalling cascade in astroglia (Porter and McCarthy, 1997), we asked if buffering intracellular Ca^2+^ with BAPTA-AM is involved, and found this to be the case (Figure 5F). Investigating this further, we blocked the calcium and calmodulin-dependent phosphatase calcineurin, which produced similar suppression (Figure 5G). However, non-selective protein kinase inhibition with the antibiotic staurosporine left the excitation-induced rise of GLT1-SEP mobility intact (Figure 5H), thus narrowing the range of the candidate molecular mechanisms involved.

## DISCUSSION

Here, we developed a functional fluorescent analogue of the main glial glutamate transporter GLT1, termed GLT1-SEP, and used it to evaluate its membrane dynamics, incorporating both surface mobility and membrane-intracellular compartment turnover, in brain astroglia. We used patch-clamp electrophysiology and super-resolution dSTORM imaging to confirm that glutamate transport properties of GLT1-SEP and its cell surface distribution on the nanoscale are fully compatible with its wild-type counterpart. Taking advantage of the pH-sensitive fluorescence and photobleaching properties of GLT1-SEP, we established that the 70-75% fraction of its cellular content reside on the astrocyte surface, with a characteristic turnover rate of 0.04-0.05 s^-1^, which corresponds to a 20-25 s cycle. That the population of functional astroglial glutamate transporters in the brain is effectively replaced several times per minute must is an intriguing discovery. Intriguingly, a recent study used fixed-tissue immunocytochemistry in pure astroglial cultures to found only ~25% of all GLT1a expressed in the cell membrane (Underhill et al., 2015), thus relating a boost in transporter numbers to the presence of neuronal connections (in the mixed cultures slices employed here).

Removing the C-terminus of GLT1-SEP only moderately increased its intracellular fraction while substantially reducing its plasma membrane lifetime. These observations suggest an important contribution of the C-terminus to the retaining of GLT1 molecules on the astrocyte surface. Intriguingly, dSTORM imaging revealed that deletion of the C-terminus severely disrupts the cell surface pattern of GLT1 and its spatial relationship with neighbouring synaptic connections (represented by clusters of PSD95).

It has previously been shown that GLT1 is endocytosed constitutively, in a clathrin-dependent manner, taking the transporter into rapidly-recycling endosomes containing EEA1 and Rab4 (Martinez-Villarreal et al., 2012). Earlier studies have also indicated that the common neuronal glutamate transporter EAAT1 also undergoes clathrin-dependent endocytosis (Gonzalez et al., 2007). Using reversible biotinylation followed by immunocytochemistry, Robinson group obtained estimates of the membrane residence time of EEAT1 (Fournier et al., 2004), and subsequent studies identified several molecular cascades that control cell surface expression of EAAT1 and GLT1 including ubiquitination and sumoylation (Gonzalez et al., 2007; Garcia-Tardon et al., 2012; Martinez-Villarreal et al., 2012; Piniella et al., 2018). While the biochemical machinery of GLT1 turnover is outside the scope of the present study, its investigation should provide further insights into the adaptive features of glutamate transport in the brain.

We next employed GLT1-SEP to investigate its lateral mobility in the plasma membrane, and the regulatory mechanisms involved. A similar question has been elegantly explored in two studies using single-particle tracking with QDs (Murphy-Royal et al., 2015; Al Awabdh et al., 2016). However, the key advantage of the present approach is that it accounts for membrane-intracellular compartment exchange, in addition to lateral mobility per se: tracking QD-labelled GLT1 must ignore the non-labelled GLT1 fraction that is being constantly delivered to the cell surface. We found relatively high average lateral diffusivity (0.10-0.15 μm^2^/s), but also a significant fraction of immobile transporters (25-30%). Importantly, the characteristic lateral diffusion time of the GLT1-SEP mobile fraction (~1.75 s) was much shorter than its membrane lifetime of (~22 s). This implies that the assessment of mobile transporter diffusivity, obtained here with GLT1-SEP or earlier with QDs, should not be noticeably influenced by its membrane turnover. Nonetheless, the latter could have a critical effect on the dynamics of the immobile (slowly moving) fraction of GLT1. For instance, the earlier studies found that GLT1 near synapses diffuse orders of magnitude slower than all transporters on average (Murphy-Royal et al., 2015; Al Awabdh et al., 2016). Thus, membrane-intracellular compartment exchange, rather than lateral diffusion, could be a preferred mechanism of the transporter turnover near synapses.

The present method has its own limitations. Similar to the QD approach, or any other live molecular tagging method, it is not technically feasible to verify fully that the labelled (or mutated) molecules have exactly the same dynamic properties as their native counterparts. Nonetheless, it is reassuring that the average lateral mobility of GLT1-SEP found here was similar to that estimated using QDs (Murphy-Royal et al., 2015), despite two very different modes of interference with the molecular structure.

The potential importance of high GLT1 diffusivity for regulating the waveform of excitatory synaptic currents was suggested earlier (Murphy-Royal et al., 2015). This might indeed be the case for large synapses, with multiple release sites (DiGregorio et al., 2002), that are prevalent in cultures or incubated slices. At small central synapses in situ, however, the kinetics of individual AMPA currents should not depend on glutamate buffering outside the synaptic cleft (Zheng et al., 2008; Savtchenko et al., 2013). Nonetheless, intense glutamatergic activity can boost glutamate escape from the cleft (Lozovaya et al., 1999), in which case lateral movement of astroglial transporters could indeed contribute to the efficiency of uptake.

Our results should provide critical real-time turnover data complementing the well-explored cellular machinery of GLT1 exocytosis and recycling in the plasma membrane (Gonzalez et al., 2007; Garcia-Tardon et al., 2012; Martinez-Villarreal et al., 2012; Piniella et al., 2018). At the same time, mechanisms that control lateral diffusion of GLT1 on the astroglial surface are only beginning to transpire. Two previous studies detected a diffusion-facilitating role of glutamate, which was either applied exogenously or released through intense neuronal network activity (Murphy-Royal et al., 2015; Al Awabdh et al., 2016), suggesting an adaptive function of GLT1 mobility. Our results confirm these observations, but also provide further important functional associations between the expected sources of molecular signalling in the brain and GLT1 mobility. We found that the deletion of the C-terminus, or the blockade of glutamate receptors, intracellular Ca^2+^ buffering, or the suppression of the calcium and calmodulin-dependent phosphatase calcineurin made the GLT1 membrane mobility irresponsive to glutamate. This is in line with previous studies which have shown that blocking kinase activity promotes glutamate uptake (Adolph et al., 2007; Li et al., 2015): lateral mobility might be one of the mechanisms assisting this process. Although regulation of GLT1 by calcineurin has previously been shown on the transcriptional level (Sompol et al., 2017), calcineurin is also known to directly dephosphorylate membrane proteins such as connexin-43 (Tence et al., 2012). At the same time, ATP application (which triggers prominent Ca^2+^-dependent cascades in astrocytes) had no effect on GLT1 mobility. We have thus identified several molecular signalling cascades that might provide important clues to the possible regulatory intervention in brain pathologies associated with malfunctioning astroglial glutamate uptake (Fontana, 2015; Peterson and Binder, 2019).

## MATERIALS AND METHODS

### DNA constructs

cDNA of rat Glt1a, cloned by Baruch Kanner group (Pines et al., 1992) under CMV promoter was a generous gift from Michael Robinson. Superecliptic pH-luorin (SEP) was introduced into second intracellular loop of GLT1a using standard cloning techniques. First, GLT1a sequence was mutated with QuikChange II Site-Directed Mutagenesis Kit [Agilent] using the following pair of primers: GTTCTGGTGGCACCTACGCGTCCATCCGAGGAG and CTCCTCGGATGGACGCGTAGGTGCCACCAGAAC in order to introduce MluI restriction site. Subsequently, SEP was amplified using pair of primers: CCGGACGCGTCTGGTTCCTCGTGGATCCGGAGGAATGAGTAAAGGAGAAGAACTTTTCAC and CCGGACGCGTTCCAGAAGTGGAACCAGATCCTCCTTTGTATAGTTCATCCATGCC ATG, which introduced linkers and enabled subcloning SEP into MluI restriction site. Resulting GLT1a-SEP was subcloned into pZac2.1 gfaABC1D-tdTomato (Addgene Plasmid #44332) (Shigetomi et al., 2013) using BmtI and XbaI sites in order to be expressed under glia-specific gfaABC1D promoter. GLT1aΔC-SEP was generated using following pair of primers: CCGATCTCGAGATGGCATCAACCGAGGGTG and CCGATGGTACCCTAGACACACTGATTAGAGTTGCTTTC which introduces “amber” stop codon after Val537 in GLT1a sequence. GLT1aΔC-SEP was then cloned to plasmid pZac2.1gfaABC1D_MCS which was generated by replacing tdTomato in pZac2.1 gfaABC1 D-tdTomato with hybridized pair of oligonucleotides: AATTCACCGGTGGCGCGCCGGATCCTGTACAACGCGTGATATCGGTACCCATATGCCGCGGACTAGTT and CTAGAACTAGTCCGCGGCATATGGGTACCGATATCACGCGTTGTACAGGATCCGGCGCGCCACCGGTG cloned into EcoRI and XbaI sites. eGFP-GLT1 was generated by amplification of GFP with the following pair of primers: CTATAGGCTAGCATGGTGAGCAAGGGCG and CGTAACTCGAGGAATTCGCCAGAACCAGCAGCGGAGCCAGCGGATCCCTTGTACAGCTCGTCCATG which introduced linker at 3’ end of GFP and enabled for it using BmtI and XhoI sites at 5’ end of GLT1a in pCMV_GLT1a plasmid. Resulting eGFP-GLT1a was subcloned into pZac2.1gfaABC1D_MCS using BmtI and XbaI restriction sites. pDisplay-SEP was generated by subcloning SEP, amplified with a pair of primers: CCGCGAAGATCTATGAGTAAAGGAGAAGAACTTTTCAC and GGCAGTCGACCTGCAGCCGCGGCCGTTTGTATAGTTCATCCATGCCATG into pDisplay-mSA-EGFP-TM (Addgene plasmid #39863) (Lim et al., 2013) using BglII and SalI restriction sites.

### Cell cultures

HEK 293T (Lenti-X 293T subclone, TaKaRa) were maintained in DMEM, high glucose, GlutaMAX (Thermo Fisher Scientific) supplemented with 10% Fetal Bovine Serum (Thermo Fisher Scientific). For transfection and patch-clamp experiments cells were plated at density 25,000 cells per 13-mm-diameter coverslip (Assistent, Germany) coated with poly-L-Lysine (Sigma-Aldrich).Cells were co-transfected with plasmids coding GLT 1 a or GLT 1 a-SEP under CMV promoter together with mRFP1 under β-actin promoter in a 2:1 ratio using Lipofectamine 2000 (Thermo Fisher Scientific) according to manufacturer instructions. Transfected cells were used for patch clamp experiments the next day.

### Electrophysiology

Patch clamp recordings were made from transfected HEK cells. Coverslips with cells were perfused with extracellular solution containing 125 mM NaCl, 2.5 mM KCl, 2 mM CaCl_2_, 1.3 mM MgSO_4_, 26 mM NaHCO_3_, 1.25 mM NaH_2_PO_4_, 12 mM D-glucose, bubbled with 95:5 O_2_/CO_2_ (pH 7.4). Patch pipettes were pulled to resistance of 4–5 MOhm when filled with the intracellular solution containing 120 mM CsCl, 8 mM NaCl, 10 mM HEPES, 0.2 mM MgCl_2_, 2 mM EGTA, 2 mM MgATP, 0.3 mM Na_3_GTP (pH 7.3). Cells were voltage-clamped at –70 mV, recordings were performed at 33°C– 35°C and signals digitized at 10 kHz. For glutamate application, we used a θ-glass pipette pulled out to an ~200 μm tip diameter, as described earlier (Sylantyev and Rusakov, 2013). Briefly, a capillary was inserted into each θ-glass channel and pressure was adjusted using the two-channel PDES-2DX-LA pneumatic microejector (npi electronic GmbH) using compressed nitrogen. θ-glass pipette was attached to Bender piezoelectric actuator (PL127.11, Physik Instrumente) and electric pulses were applied via a constant-voltage stimulus isolator (DS2, Digitimer).

### Primary dissociated culture

Dissociated hippocampal cultures from P0 (postnatal day 0) Sprague-Dawley rats were prepared in full compliance with the national guideline and the European Communities Council Directive 0f November 1986, and the European Directive 2010/63/EU on the Protection of Animals used for Scientific Purposes. Brains were removed and hippocampi were isolated on ice in dissociation medium - DM (81.8 mM Na_2_SO_4_, 30 mM K_2_SO_4_, 5.8 mM MgCl_2_, 0.25 mM CaCl_2_, 1 mM HEPES pH 7.4, 20 mM glucose, 1 mM kynureic acid, 0.001% Phenol Red), hippocampi were later incubated twice for 15 minutes at 37°C with 100 units of papain (Worthington, NY) in DM and rinsed three times in DM and subsequently three times in plating medium (MEM, 10% fetal bovine serum (FBS) and 1% penicilin-streptomycin; Thermo Fisher Scientific). Hippocampi were triturated in plating medium until no clumps were visible and cells were diluted 1:10 in OptiMEM (Thermo Fisher Scientific), centrifuged for 10 minutes at room temperature, at 200 x g. The resulting cell pellet was suspended in plating medium, cells were counted in 1:1 dilution of 0.4% Tryptan Blue solution (Thermo Fisher Scientific) and plated at density 75,000 cells per 13-mm-diameter coverslip (Assistent, Germany) coated with 1 mg/ml poly-DL-lysine (Sigma-Aldrich, P9011) and 2.5 μg/ml laminin (Sigma-Aldrich, L2020). Three hours after plating medium was exchanged for maintenance medium (Neurobasal-A without Phenol Red, 2% B-27 supplement, 1% penicillin-streptomycin, 0.5 mM glutaMAX, 25 μM β-mercaptoethanol; ThermoFisher Scientific) and cells were kept at 37°C, under a humidified 5% CO_2_ atmosphere. Cells were transfected with plasmids using Lipofectamine 3000 (Thermo Fisher Scientific) at 7-10 days in vitro (DIV). Lipofectamine – DNA complexes were prepared according to manufacturer’s instructions and were incubated with cells for 1 h in the incubator, in fresh transfection medium (MEM without Phenol Red, 2% B27 supplement, 1mM pyruvate, 0.5 mM GlutaMAX, 25 μM β-mercaptoethanol; Thermo Fisher Scientific). After transfection conditioned maintenance medium was returned to cells. All experiments were performed at 14–19 DIV.

### Organotypic hippocampal culture

Transverse hippocampal organotypic cultures were prepared according to Stoppini and colleagues (Stoppini et al., 1991) with some modifications. P8 Sprague-Dawley rats were sacrificed in full compliance with the national guideline and the European Communities Council Directive of November 1986, and the European Directive 2010/63/EU on the Protection of Animals used for Scientific Purposes. Hippocampi were dissected in ice-cold Gey’s Balanced Salt Solution (Merck) supplemented with 28 mM glucose, 1 mM Kynureic acid and 10 mM MgCl_2_, and 350 μm hippocampal slices were cut using McIlwain tissue chopper. Slices were cultured on 0.4 μm Millicell membrane inserts (Merck) in Minimum Essential Medium (MP Biomedicals) supplemented with 25% Hank’s Balanced Salt Solution (MP Biomedicals), 25% horse serum, 1% Penicillin-Streptomycin, 1 mM GlutaMax (all Thermo Fisher Scientific), and 28 mM Glucose (Sigma-Aldrich). Medium was changed 3 times per week. After 4 DIV, cultures were transfected with plasmids using a biolistic method (Helios Gene Gun, Bio-Rad). To obtain sparse astrocyte labelling we used 1 μm gold particles (Bio-Rad) and followed a standard protocol (Benediktsson et al., 2005) for preparation of gene gun bullets. Slices were shot at 160 PSI Helium pressure using modified gene gun barrel, in accord with accepted routines (Woods and Zito, 2008), where diffuser screen were replaced with stainless steel wire mesh (180 mesh per inch, 36% open area; Advent Research Materials Ltd.). Slices were used for experiments 4-10 days after transfection.

### Imaging and FRAP

Imaging was performed using an Olympus FV1000 system under Olympus XLPlan N25 x water immersion objective (NA 1.05). Imaging system was linked to two mode-locked, femtosecond-pulse Ti:Sapphire lasers (MaiTai from SpectraPhysics-Newport and Chameleon from Coherent), first one for imaging, was set at a wavelength of 910 nm and the other was for bleaching set on 690 nm, each of the lasers was connected to the microscope via an independent scan head. 690 nm for bleaching was selected based on the 2-P excitation spectrum for GFP (Drobizhev et al., 2011). The imaging laser power was kept below 4 mW under the objective at all times to minimize phototoxic damage, a power range validated by us previously in similar settings (Jensen et al., 2019). Bleaching laser power was kept around 10 mW. Dissociated mixed cultures were imaged in extracellular solution containing: 125 mM NaCl, 2.5 mM KCl, 30 mM Glucose, 25 mM HEPES, 2 mM CaCl_2_ and 1.3 mM MgSO_4_; pH 7.4, at 32-34°C. In puffing experiments, pH 5.5 extracellular solution contained: 125 mM NaCl, 2.5 mM KCl, 30 mM Glucose, 25 mM MES, 2 mM CaCl_2_ and 1.3 mM MgSO_4_, Extracellular solution with 50mM NH_4_Cl contained: 50 mM NH_4_Cl, 75 mM NaCl, 2.5 mM KCl, 30 mM Glucose, 25 mM HEPES, 2 mM CaCl_2_ and 1.3 mM MgSO_4_, pH 7.4.

Organotypic cultures were imaged in artificial cerebrospinal fluid (aCSF) containing: 125 mM NaCl, 2.5 mM KCl, 2 mM CaCl_2_, 1.3 mM MgSO_4_, 26 mM NaHCO_3_, 1.25 mM NaH_2_PO_4_, 20 mM D-glucose, 0.2 mM Trolox, bubbled with 95:5 O_2_/CO_2_ (pH 7.4) at 32-34°C.

FRAP experiments were performed using 22 x zoom at 256 x 256 numerical resolution, resulting in a ~0.09 μm pixel size. Frame size was kept constant: 138 x 80 pixels, giving 148.32 ms per frame (unidirectional scanning, 4.0 μs pixel dwell time). Bleached region was kept constant – a 18-pixels diameter circle (2.06 μm^2^) scanned with second laser using tornado mode, resulting in fast bleaching time – 46 ms. In some experiments drugs (1 mM Glutamate or 100 μM ATP) were puffed for 250 ms just before the bleaching using Pneumatic PicoPump (World Precision Instruments). Imaging, bleaching and puffing were synchronized using Axon Digidata digitizer (Molecular Devices).

For whole cell FRAP (only in dissociated culture) 512 x 512 pixel frames were imaged every 1.644 s. Bleached region – 398-pixels diameter circle was scanned with second laser using tornado mode resulting in fast bleaching time – 2.00 s. In order to image and bleach as big astrocyte surface as possible, we used 2 to 5 x zoom resulting in corresponding pixel size 0.497 μm to 0.198 μm. Pixel size was taken into account for data analysis and calculations.

In some FRAP experiments (see Results) we used the following drugs in the bath solution: TTX (1 μM, Tocris), MPEP (1 μM, Tocris), NBQX (10 μM, Tocris), LY 341495 (30 nM, Tocris), YM (300 nM, Tocris), Bicuculine (10 μM, Sigma-Aldrich), 4-AP (4-Aminopyridine, 500 μM, Sigma-Aldrich), FK-506 (1 μM, Sigma-Aldrich), Staurosporine (100 nM, Cell Signaling Techn.).

### Super-resolution microscopy

We used the single-molecule localization microscopy (SMLM) technique direct stochastic optical reconstruction microscopy (dSTORM) (van de Linde et al., 2011; Endesfelder and Heilemann, 2015) as described previously (Heller et al., 2017; Heller and Rusakov, 2019; Heller et al., 2020). Naïve dissociated hippocampal cultures and cultures expressing either GLT1a-SEP or GLT1aΔC-SEP were fixed using 37°C prewarmed 4% paraformaldehyde in PEM buffer (80 mM PIPES pH 6.8, 5 mM EGTA, 2 mM MgCl_2_) (Leyton-Puig et al., 2016; Pereira et al., 2019) for 10 minutes at 37°C. Then, cells were washed thrice in PBS, incubated in 0.1% NaBH_4_ in PBS for 7 minutes, washed thrice with PBS and incubated in 10 mM CuSO_4_ in 50 mM NH_4_Cl, final pH = 5 for 10 minutes. Cells were washed thrice with water quickly and once with PBS. Cells were then permeabilised and blocked with PBS-S (0.2% saponin in PBS) supplemented with 3% BSA for 1 hour. Afterwards, cells were incubated with primary antibody (see below) in PBS-S overnight at 4°C, washed trice with PBS-S, incubated with secondary antibody (see below) in PBS-S for 2 hours, washed twice with PBS-S and twice with PBS. Lastly, cells were post-fixed with 4% paraformaldehyde in PBS, washed thrice with PBS and stored at 4°C until being prepared for imaging.

Primary antibodies used: post-synaptic protein PSD-95 (mouse, 6G6-1C9, recombinant rat PSD-95, Novus Biologicals, NB110-61643, AB_965165, dilution 1:500), glial glutamate transporter GLT1 (guinea pig, polyclonal, synthetic peptide from the C-terminus of rat GLT1, Merck, AB1783, AB_90949, dilution 1:1,000), GFP (chicken, polyclonal, GFP directly from *Aequorea Victoria*, Thermo Fisher Scientific, A10262, AB_2534023, dilution 1:1,000).

Secondary antibodies used: anti-mouse IgG (donkey, CF568-conjugated, Biotium, 20105, AB_10557030, dilution 1:500), anti-chicken IgY (goat, Alexa647-conjugated, Thermo Fisher Scientific, A21449, AB_1500594, dilution: 1:1,000), anti-guinea pig IgG (donkey, Alexa647-conjugated, Jackson ImmunoResearch Labs, 706-606-148, AB_2340477, dilution: 1:1,000).

Images were recorded with a Vutara 350 microscope (Bruker) in photo-switching buffer containing 100 mM cysteamine and oxygen scavengers (glucose oxidase and catalase) (Metcalf et al., 2013). Images were recorded with frame rate of 33 Hz (561 nm for CF568) or 66 Hz (640 nm for Alexa647). Total number of frames acquired per channel ranged from 3,000 to 20,000. Data were analysed using the Vutara SRX software (version 6.02.05). Fiducial markers (100 nm TetraSpeck microspheres, T7279, Thermo Fisher Scientific) were used for drift correction.

### Cluster and Nearest-neighbour analysis

In dSTORM maps, clusters of PSD95 were identified using DBScan, a well-established density-based clustering algorithm (Ester et al., 1996), with a minimum of 50 particles per cluster and a maximum particle distance of 100 nm; the latter parameters correspond to 250-300 nm wide PSD5 clusters which are consistent with the typical PSD size at common central synapses (Chen et al., 2008). The distribution of nearest-neighbour distances *D*(*r*) between PSD95 clusters and GLT1 molecular species (and also among GLT1 molecular species) was calculated as the occurrence of distances *r*, with a 5 nm or 10 nm binning step, and normalised to the overall number of registered events. To assess non-uniformity of the experimental distribution pattern *D*(*r*) was compared to the theoretical *D*(*r*) of a 2D Poisson point process (evenly random scatter) of the same surface density *λ*, in the form *D*(*r*) = 1 – exp(−*λπr*^2^) (Stoyan, 2006).

### FRAP data analysis

Raw images were analysed using ImageJ. Mean fluorescence intensity was calculated for manually selected ROIs: Background - manually selected background ROI outside of transfected cell (FBKG), reference ROI which was manually outlined transfected cell in the imaged frame (FREF). For each frame mean fluorescence intensity of bleached ROI (FBL) was normalized according to the formula: F_NOR= (F_BL-F_BKG)/(F_REF-F_BKG). Normalized fluorescence value at the frame after the bleaching pulse (close to the background value) was subtracted from all values in data set. Finally, resulting fluorescence values were normalized to 40 frames before bleaching.

For the whole-cell bleaching experiments we performed similar analysis however for each cell we have measured mean fluorescence in manually selected 3 ROIs (FROI) defined as ~10 μm diameter circle placed outside of the cell soma. Additionally for each analysed data set we measured mean fluorescence in manually selected background ROI (FBKG) – outside of transfected cell and in reference ROI (FREF) which was manually outlined transfected cell outside of bleached region. We performed the same normalization for each specific ROI as described above. The kinetic analyses of membrane turnover and FRAP traces were carried out as detailed in Figs S1–S2. The FRAP time course 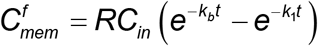 (notations in Figure 1-figure supplement 1–S2) was fitted using the non-linear fitting routines ExpGroDec (exponent fitting) in Origin (OriginLab).

To evaluate lateral diffusivity from the spot-FRAP kinetics, we used the well-established Soumpasis method for circular ROIs (Soumpasis, 1983; Kang et al., 2009), in which the fluorescence time course is fitted with the equation 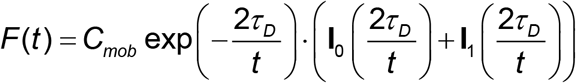 where *C_mob_* is the mobile fraction, 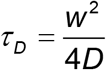, *w* is ROI radius, *D* is diffusion coefficient, and **I**_0_ and **I**_1_ are modified Bessel functions of the first kind; this fitting has only two free parameters, *C_mob_* and *D*. The fitting was carried out using Soumpasis in the Origin software (OriginLab). The average diffusivity *D*\* was therefore calculated as *D*\* = *C_mob_ · D*.

Statistical inference was calculated using Origin’s Hypothesis Testing from individual ROIs as statistical units (2-4 per cell): routine two-way ANOVA tests indicated no significant influence of the cell identity factor on the effects of experimental manipulations under study.

## AKNOWLEDGEMENTS

The study was supported by the Wellcome Trust Principal Fellowship (212251_Z_18_Z), ERC Advanced Grant (323113) and European Commission NEUROTWIN grant (857562), to DAR.

## AUTHOR CONTRIBUTIONS

DAR and PM narrated the study; PM designed fluorescent GLT1-SEP and carried out FRAP experiments and data analyses; JH designed and carried out dSTORM analyses; DAR designed experiments and carried our data analyses; DAR wrote the manuscript, which was contributed to by PM and JH.

